# What rhythmic perception and amusia can tell us about vocal social communication in schizophrenia

**DOI:** 10.1101/092098

**Authors:** Rachel L. C. Mitchell, Krystal Gamez, Joshua Bolgar, Eli S. Neustadter, Monica E. Calkins, Tyler M. Moore, David I. Leitman

## Abstract

**Background:** Perceiving social intent throughvocal intonation is impaired in schizophrenia; thisdysprosodia partly arisingfrom impaired pitch perception.Individuals with amusia (tone-deafness) are insensitive to pitch change andalso demonstrate prosody deficits. Sensitivity to rhythm is reduced in amusia when tonal sequences contain pitch changes (polytonic), but is normal for monotonic sequences, suggesting perceptual impairment originates at a secondary processing stage where pitch- and time-relatedcues are yoked. Here, we sought to ascertain: 1) whether schizophreniapatients demonstrate rhythmic deficits, 2) whether suchdeficits are restricted to polytonic sequences, and 3) how pitch and rhythm perception relate to prosodic processing.

**Methods:** Seventy-sixparticipants (33 schizophrenia) completed tasks assessing pitch and prosody perception, as well as monotonic and polytonic rhythmic perception.

**Results:** Increasing tone-deafness correlated with pitch-dependent rhythm detection impairments. Pitch and prosody correlated across all participants. Schizophreniapatients displayed basic time and pitch deficits. Correlations and path analyses indicated prosodic processing is an associatedfunction of pitch and pitch-dependent rhythm perception,with pure temporal processing playing an indirect role.In schizophrenia, deficits in monotonic and polytonic rhythmic perception did not contribute to prosodic processing dysfunction, and montonic rhythmic dysfunction and pitch perception did not covary.

**Conclusions:** Exploring similarities between amusia and schizophrenia focused our characterization of prosodic processing as the function of sub-processes reflecting pitch and time perception,whichare prerequisite for prosodic processing. The uniqueness of dysprosodia in schizophrenia relative to other illnesses may be measured by idiosyncrasy in the pattern and magnitude of the sub-process task relationships.

## INTRODUCTION

Schizophreniais a neuropsychiatric disorder that affects cognitive, social, and affective functioning. Cardinal among these impairments isdifficulty in perceiving social intent via vocalintonation(prosody)(1–3). A number of recentstudies have reconceptualizeddysprosodia in schizophrenia from being an experiential affective deficit, to being a perceptual-cognitive one, linking dysprosodia to basic dysfunction in primary auditory perceptual systems and the ability to discriminatepitch(2;4). Pitch is a cardinal cue for perceiving emotion(5;6), and inabilities in its perceptionplay a prominent role in schizophrenia dysprosodia(7).

The pattern of these pitch deficits closely parallelsthose shown in the neurodevelopmental disorderof congenital amusia. Amusia is a disorder that affects roughly 4% of the population worldwide (8). It is characterized by profound pitch insensitivity that impairs music processing skills as well as short term memory of pitch and timbre information,whilst leaving other auditory perceptions and languagefunctions largely intact(9–11).Such tone-deafness likely has strong genetic roots, as it displays a clearMendelian inheritance pattern(9;12;13).

Similarities between the sequelae of tone deafness in amusiaand schizophrenia extend to impairments in the perception of emotional prosody and the use of fundamental frequency (F_0_percieived pitch) as an affectively informative cue (2;3;6;14;15), as well as the utilization ofterminal pitch changes in perceiving interrogative versus declarative intent (16–18). Other similarities include electrophysiological abnormalities in automatic pitch and duration deviation recognition using mismatch negativities (MMN)(19–23). Moreover,electrophysiological and neuroimaging data from people withamusiaduring the processing of pitch intonation changes demonstrate a pattern of dysfunctional activity and coordination between superior temporal and inferior temporal gyrithat is also observed during the processing of prosody in schizophrenia(3;24–26). Finally, in both amusia and schizophrenia, prosodic processing deficits also correlate with structural abnormalitiesin the dorsal auditory pathway from Wernicke's area to Broca's area(18;27). Additionally,non-congenital amusia,though rare,typically results from restricted lesions in the right frontal cortex(28;29).

In addition to spectral cues like pitch,otherprincipal cuesfor the perception ofprosody are temporal cues such asmeter or rhythm(5;6). To date, the perception of rhythm in schizophrenia has not been studieddirectly. What is known though is that patients with schizophrenia display a wide variety of auditory processing deficits in response to changes in the physical properties of the auditory environment. Such changes include deficits relating to stimulus pitch, intensity, and duration,as evidenced by MMN and p300 oddball paradigms(22;23;30). A deficit in temporal duration perceptionmight well extendto an impaired ability to perceive rhythm.

In amusia, Hyde and Peretz(31)and Foxton et al. (32) revealed rhythmic deficits only in tasks in which the pitch of tonal sequences also varied. These results were surprising, given that conventional wisdom buttressed by lesion data, supported the notion of doubly-dissociated processing systems for rhythm and pitch(29;33). Conversely, other studies such as that ofBoltz(34)indicated that healthy individuals fair poorly on musical judgments of melodies in which the pitch and rhythmic patterns conflict.To account for this divergent evidence,Hyde and Peretz(31)and Foxton et al.(32) have proposed a two-tiered network in which early and independent processing of pitch and rhythm are yoked together at some later stage.

In the current study, given the similarities between schizophrenia and amusia in the processingof music and prosody, we sought to explore the following questions:1) Would schizophrenia patients’ rhythmic performance vary in the presence of tonal changes in rhythmic sequences? 2) Would such rhythmic performance variations themselves vary according to the acuity of a participant’s pitch perception?3) Does rhythmic performance contribute to dysprosodia in schizophrenia? We hypothesized that patients with schizophrenia would display rhythmic deficits in pure meter perception, as well as deficits that were separately dependent on pitch, and that these deficits would contribute to prosody perception deficits in thisdisorder.

## METHODSAND MATERIALS

### Participants

Thirty-threeindividuals with schizophrenia and forty-fourhealthy controlparticipants with no prior history of mental illness participated in this study. As indicated in **Table 1**, age, gender, and parental education were comparable across the two groups. Participantshad no history of hearing loss as measured by the American Academy of Otolaryngology-Head and Neck Surgery’ 5-minute hearing test questionnaire;however,one controlparticipant was excluded from the final dataset because of hearing deficits identified during the test session. Informed consent was obtained from all participants and the University of Pennsylvania’s Institutional Review Board approved all procedures.

**Table 1:** Demographics, clinical symptoms and ancillary neuropsychological measures.

|  | **Healthy Controls**<br>**(N=43)** | **Schizophrenia**<br>**(N=33)** |
| --- | --- | --- |
| Age (years) | 34.2±12.4 | 35.5±9.5 |
| Gender (#female) | 17 | 11 |
| Handedness R (L) | 43(0) | 33(0) |
| Average parental education (years) | 14.1±2.5 | 13.1±2.5 |
| Medication (CPZ equiv dose) | NA | 483.8±396.9 |
| *Negative symptoms (SANS)<sup>a</sup>* |  |  |
| Total global score | NA | 9.6±4.2 |
| Affective flattening | NA | 2.4±1.2 |
| Alogia | NA | 1.3±1.3 |
| Avolition | NA | 1.9±1.2 |
| Anhedonia | NA | 2.9±1.2 |
| Attention | NA | 1.1±1.5 |
| *Positive symptoms (SAPS)<sup>a</sup>* |  |  |
| Total global score | NA | 5.1±3.6 |
| Hallucinations | NA | 1.8±2.0 |
| Delusions | NA | 2.3±1.7 |
| Bizarre Behavior | NA | 0.5±0.9 |
| Formal thought disorder | NA | 0.5±1.1 |
| *Ancillary Neuropsychological Measures:* |  |  |
| Tone Matching (Pitch)(% correct)<sup>b</sup> | 0.79±0.12 | 0.69±0.08 |
| Prosody (% correct)<sup>c</sup> | 0.64±0.11 | 0.55±0.12 |
| Rhythm – Monotonic Threshold Log<sup>d</sup> | 3.11±0.96 | 3.96±0.92 |
| Rhythm – Polytonic Threshold Log<sup>e</sup> | 3.44±1.1 | 4.36±1.23 |

Caption: All values represent mean ± standard deviation. <sup>a</sup>SANS/SAPS ratings collected on all patients.
<sup>b</sup>( $t_{72}= 4.24$ , $p<0.01$ ); <sup>c</sup>( $t_{63}= 3.35$ , $p<0.01$ ); <sup>d</sup>( $t_{66}= -3.83$ , $p<0.01$ ); <sup>e</sup>( $t_{62}= -3.28$ , $p<0.01$ );

### Procedures

#### Pitch Task

Pitch perception was assessed using the same Tone Matching Taskas used in a number of previous studies (2;22;35)Participants were presented with pairs of 100ms tones with 500ms inter-tone intervals. To avoid learning effects, 3 different base tones (500, 1000, and 2000 Hz) were utilized, to representthe frequencies considered to be particularly important for conveying speech (36). Of the 94 pairs of stimuli, half were identical, and half differed in frequency by 1%, 1.5%,2.5%, 5%, 10%, 20%, or 50% of their counterpart. Participants were asked to indicate whether the pitch of the two tones was the same or different. One participant(a patient with schizophrenia) responded identicallytoevery item and was thus excluded from the analyses of data forthis task.The output variable used in the subsequent data analyses was the percentage of tone matching trials correctly identified as same or different.

#### Rhythm Task

The rhythm tasks used in this study mirroredthe asynchronous rhythm task employed by Foxton et al.(32). Two tasks, a monotonic- and a polytonic-version, were designed to measure rhythm perception. In both tasks, participants listened to two rhythmic sequences of tones and were asked to identify whether it was the first or second rhythm that contained an additionalgap. Each rhythm compriseda series of five 100ms tones with a standard inter-tone onset interval of 300ms. The increase in variable gap across one interval (aperiodicity)that participants were asked to detect began at 256ms, and increased or decreased adaptively based on performance in a two-down, one-up procedure that required participants to correctly identifytwo trialsto decrease the length of the gap, which made the task harder, but required only one trialto be answered incorrectly to increase the length of the gap, which made the task easier. Adaptive change increased or decreased the gap first by 128ms, then by 64ms, 32ms, 16ms, and 8ms, and finished with 5 repetitions of 4ms, for a minimum of 10 reversals or 22 trials. There was a 1.5s gap between the two rhythms in each item. The tasks were identical except for the frequencies of the tones used. In the monotonic task, all tones were presented at 1000Hz. In the polytonic version of the task, tones were presented in random pitches taken from a series of notes derived from an octave divided into seven steps (828.1, 914.3, 1009.4, 1114.5, 1230.5, 1358.6, and 1500Hz). Participants with rhythm thresholds more than 2.5 standard deviations above the mean (1 control and 2patients with schizophrenia for both the monotonic and polytonic tasks) were winsorized to their closest neighbor below this threshold.The thresholdlevels used in the subsequent data analyses for each rhythm task were calculated by averaging the lengthened intervals at the last five points of the difficulty-level alternations.

#### Prosody Task

Prosody perceptionwas assessed usinga subset of well-validated stimuli taken from Juslin and Laukka(6), that featured British English speakers instructed to state neutral sentences (e.g. “it is 11 o’clock”) whilstprojecting a specifc emotion. The emotions covered in our taskincluded anger, fear, happiness, and sadness, as well as a neutral ('no expression’) condition. Each emotion was presented 6 times, while given that large variation in pitch is most classically predictive of happiness, this emotionwas presented 9 times, to assist in the detection of the relationship with pitch perception. The task thus totalled 33 trials(37;38). From these forced-choice response options, participants were asked to choose which emotion was represented by intonation. The output variable used in the subsequent data analyses was the percentage of trials for which the conveyed emotion was correctly identified. Five participants completed a similar prosody task within a different experimental protocol and were thus excluded from the analysis of this task.

#### Clinical symptoms and sociality assessments

Positive and negative clinical symptoms were measured using the SANS (Scale for the Assessment of Negative Symptoms(39)) and SAPS (Scale for the Assessment of Positive Symptoms(40)).

### StatisticalAnalysis

Analysis of rhythmic task performance was conducted in the following manner. Rhythmic thresholds (given in milliseconds) were found to be highly non-normally distributed, hence these values were log-transformed for statistical analysis. Multivariate analysis was applied torhythmic performance with a within groups factor for task (monotonic and polytonic) and a between groups factor of diagnosis (controlversusschizophrenia). In order to directly assess the influence of pitch perception on rhythmicity judgements, a second across-diagnosis multivariate analysis was performed in which a group factor contrasting low, intermediate, and high levels of pitch acuity based on the normalized performance on the tone matchingtask was substituted for diagnosis.

Assessment of the relationship between pitch perception, affective prosody, and rhythm perceptionwas implementedusingPearson correlations, followed by confirmatory path analysis to specifically examine the roles of pitch and rhythm (polytonic and monotonic) task performance on prosodic processing within and across groups. Our initial fully-connected hypothetical model (see **Figure 2a** & caption) was fully saturated, therefore, we iteratively pruned the model paths. Model integrity was assessedusing root mean-square error of approximation (RMSEA),comparative fit index (CFI),and standardized root mean-square residual (SRMR).Structural equation modeling was conducted using Mplus V7 software (Muthén & Muthén,www.StatModel.com). Correlations between clinical, sociality assessment and task performance employed Spearman coefficients (R_*s*_) when outliers, as seen in scatterplots, deemed it warranted. Excluding structural equation modeling, allstatistical tests were conducted in R (http://www.r-project.org),employing an α criterion of <0.05.

## RESULTS

### Rhythmic perception in schizophrenia

Patients’ rhythmic perception was significantly impaired (F_1,72_= 14.34, p<0.01). Overall, participants also performed better on the monotonic than the polytonicrhythm task, (F_1,70_= 10.04, p<0.01). The interaction between diagnosis and rhythm task was not significant(F_1,70_= 0.17, p> 0.68).

### Rhythmic perception as a function of pitch acuity

Scatterplots of pitch by prosody task performance indicated that schizophrenia and controls participants performed along a continuum; with patients predominating the lower ends of pitch-prosody performance and controls the higher levels of pitch-prosody perfromance (**Figure 1A**). Therefore, all participantswere placed into groups based on performance on the Tone Matching Taskthat assessed pitch perception(**Figure 1B**), rather than according to diagnosis. Scores on the Tone Matching Task were translated into zscores.Participantswith a z score of 0.5 or above (n=25) were classified into the High performance group. Participants scoring between -0.5 and 0.5 (n=25) were classified into an Intermediate performance group. Finally, participants with a z score of -0.5 or below (n=25) were classified into the Low performance group. Overall, rhythmic thresholds varied across High, Intermediate and Low pitch groups(F_2,66_ = 9.68, p<0.01). Polytonic rhythmthresholds were again significantly higher than monotonicthresholds (F_1,66_= 8.24, p<0.01). Importantly, contrasting monotonicand polytonic performanceas a function of pitch group using protected Tukey contrasts indicated that polytonic rhythmthresholds increased in reference to monotonicthresholdsfor the low vs. high, and the intermediate vs. high Tone Matching Task groups(p< 0.01 and p< 0.01respectively), but not between the intermediate vs. low groups(p>0.7), this latter contrast leading to a non-significant pitch group by task level interaction(p>0.4).

**Figure 1.**
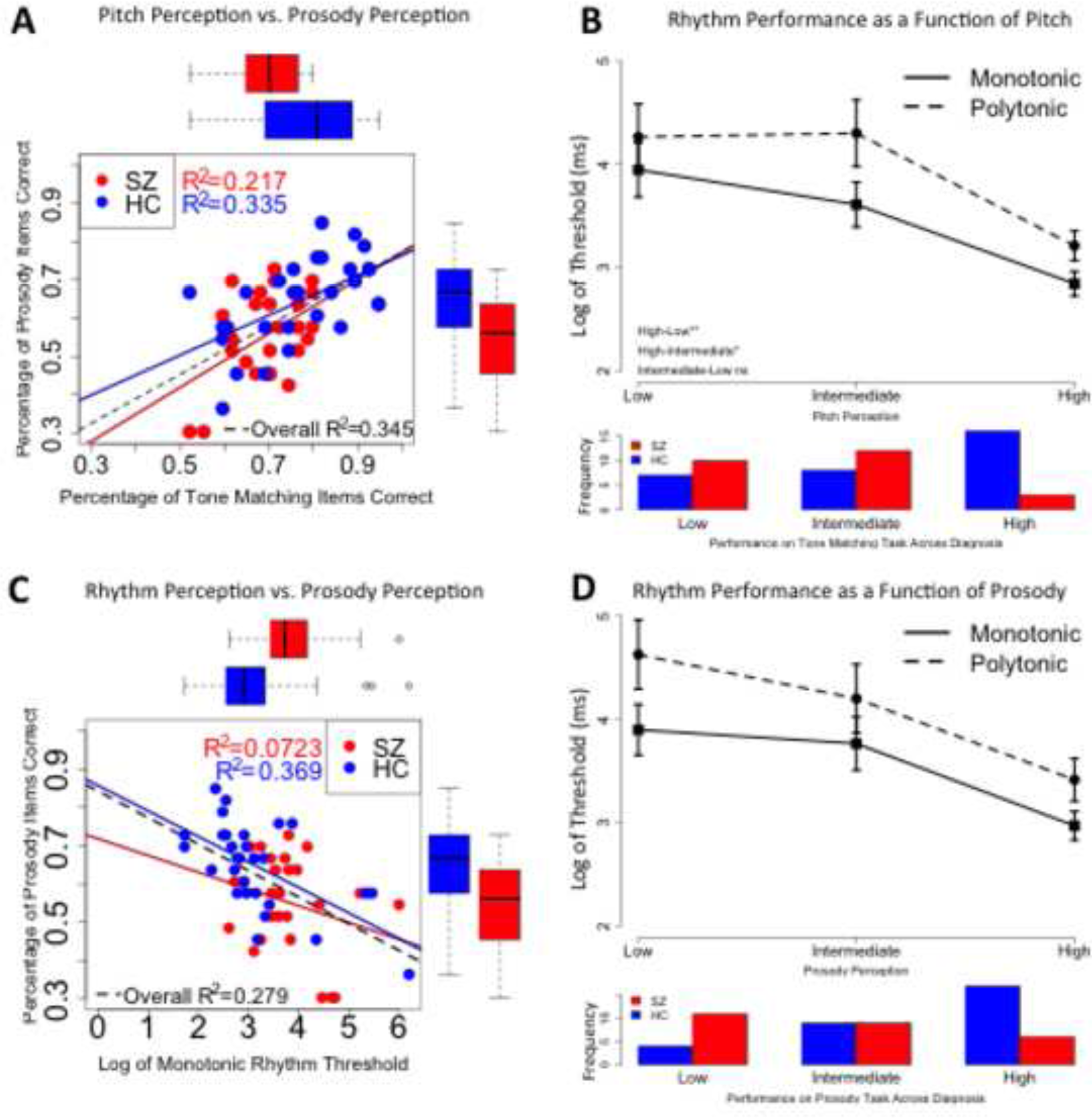
Comparisons of performance on pitch, prosody, and rhythm tasks across control participants (HC) and schizophrenia participants (SZ). **(A)**Pitch and prosody performance comparison indicates significant correlations between and across groups. X and Y box & whisker (top and right) graphs display group differences in pitch and prosody. **(B)** *Top:* Monotonic and polytonic rhythm performance across pitch acuity groups. Results indicate significantly higher detection thresholds for polytonic vs. monotonic for high and both intermediate and low pitch acuity. *Bottom*: Distribution indicates the control participants dominate high pitch acuity group *x*^2^ =9.69, p<0.01. **(C)** Monotonic rhythm and prosody comparison indicates significant correlations only in healthy control group. X and Y box & whisker (top and right) graphs display group differences in rhythm and prosody. **(D)**Top: Monotonic and polytonic rhythm performance across prosodic acuity groups. Again, results indicate significantly higher detection thresholds for polytonic vs. monotonic for high and both intermediate and low pitch acuity. *Bottom*: Distribution indicates the control participants dominate high prosodic acuity group *x*^2^ =8.28, p<0.02.

### Rhythmic and pitch acuityand their contributions to prosody

As expected, control participants significantly outperformed schizophrenia participants on both the pitch(t_72_= 4.25, p<0.01) and prosody (t_63_= 3.35, p<0.01) tasks. Correlations between tasks indicated robust correlations within patients (r_29_= 0.54, p<0.01) and controls (r_37_= 0.46, p<0.01) and across groups for pitch and prosody (r_68_= 0.55, p<0.01)(**Figure 1A**).Across groups (r_67_= -0.50, p<0.01) and within the healthy controls(r_36_= -0.58, p<0.01),monotonic rhythm thresholds correlated with prosody task performance. Yet this correlation was not observed in the patients withschizophrenia(r_28_= -0.24, p>0.19) (**Figure 1C**).In both theschizophrenia (r_28_= 0.37, p<0.05) and control groups(r_41_= 0.77, p<0.01), polytonicand monotonic thresholdswere correlated. To contrast the relative acuity of polytonic rhythmversus monotonic performance as a function of prosody,we employed the same analysis that we applied to pitch in **Figure 1B**. We z-transformed each participant’s prosodytask performance, and created high (n=29), intermediate (n=24), and low performing groups (n=18), finding a nearly identical pattern of rhythmictask performance to that observed for the pitch task performance groups (**Figure 1D**): Overall, rhythmic thresholds varied significantly across thehigh, intermediate and low prosody task performance groups (F_2,63_= 8.79, p<0.01). Strikingly, as was the case with the pitch versus rhythm analysis, contrasting monotonic and polytonic rhythm performanceas a function of prosodytask performance group indicated that polytonic thresholds increased relative to monotonicthresholds across low to high and intermediate to high prosody task performancegroups (p<0.01 and p<0.01, respectively), but not between intermediate and low groups(p>0.80), this latter contrast again leading to a non-significant prosody group by rhythm type interaction(p>0.61).

The pattern of pairwise task correlations(**Table 2**)fully delineated the structure and relative magnitude and manner of time (i.e. rhythm) and pitch perception to prosodic functioning. Therefore, we employed our task covariance matrix to test a path model of prosodic processing based on our *a priori* hypothesis. **Figure 2** describes this model as applied within the controland schizophreniagroups. Removal of themonotonic rhythmto prosodypath yielded a robust non-saturated model within controls. This model was less than robust within schizophrenia alone, which may be due to its smaller sample size. Whereas within control participants, pitch-monotonic rhythmcovariance was robust (β=-0.62,p<0.0001), within schizophrenia, itwas not significant (β=-0.08, p=0.651). Additionallytherobust monotonic rhythm-prosodypath present in controls (β=0.75, p<0.0001) was substantially weakerinschizophrenia(β=0.39, p=0.01; see **Figure 2**).Overall, these models indicated that pitchrobustly contributed to prosodydirectly. In contrast,monotonic rhythm, while not directly contributing to prosody, rather robustly contributed to polytonic rhythm and thus indirectly to prosodyvia the polytonic rhythm-prosody path.

**Figure 2.**
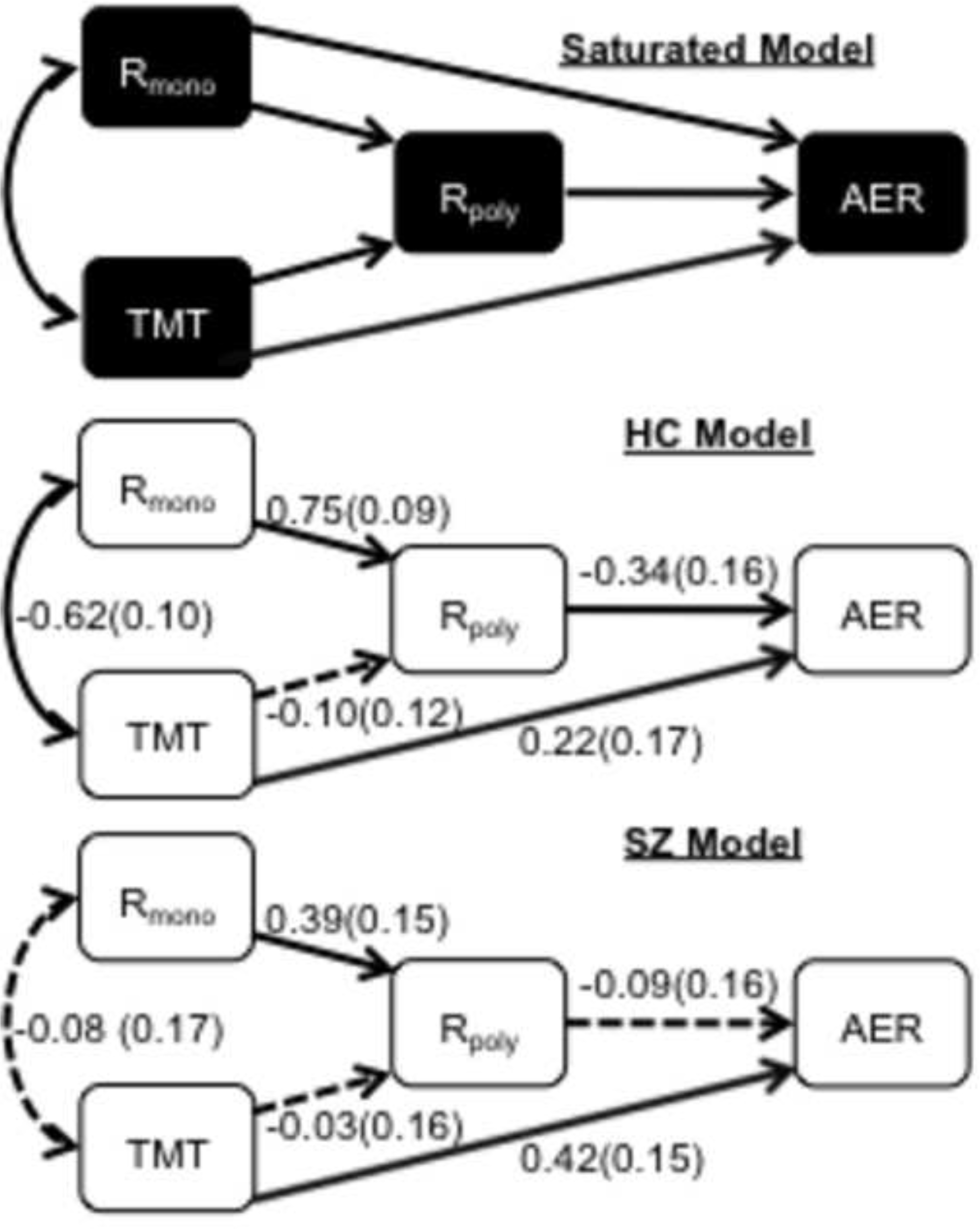
Confirmatory path analysis between monotonic rhythm (R_mono)_), polytonic rhythm (R_poly_), pitch performance (TMT) and prosody performance (AER) across groups and between control participants (HC) and schizophrenia participants (SZ). **(A)** Our initial hypothetical fully connected model was saturated. **(B)** In healthy participants, removal of the R_mono_-AER path yielded the best fit model and was hence adopted. Solid lines indicate significant (p<0.05) standardized path coefficients (β). There is strong TMT-R_mono_ covariance, direct contributions from TMT to AER,and significant mediated contributions to AER from rhythmic performance, however, no-significant TMT- R_poly_ path(dashed line)(*x*^2^=0.22, p=0.64; RMSEA<0.001, CI± 0.32, probability of RMSEA ≤0.05=0.66; CFI=1.00; SRMR=0.01) (C) In contrast to the healthy participant model, the schizophrenia group model displayed no-significant TMT-R_mono_ covariance and a non significant overall R_poly_–AER path.Overall model fit in the schizophrenia model was less robust than that found within healthy controls (*x*^2^=4.26, p=0.04; RMSEA=0.31, CI± 0.32, probability of RMSEA ≤0.05=0.05; CFI=0.71; SRMR=0.08).

**Table 2:** Pairwise task correlations of monotonic rhythm (R_mono)_), polytonic rhythm (R_poly_), pitch performance (TMT) and prosody performance (AER) across groups and between healthy control participants (HC) and schizophreniaparticipants (SZ).

| **ACROSS GROUPS** | $R_{\text{mono}}$ | $R_{\text{poly}}$ | TMT |
| --- | --- | --- | --- |
| $R_{\text{mono}}$ | | | |
| $R_{\text{poly}}$ | 0.65 | | |
| TMT | -0.54 | -0.50 |  |
| AER | -0.50 | -0.39 | 0.55 |
| **HC** | $R_{\text{mono}}$ | $R_{\text{poly}}$ | TMT |
| $R_{\text{mono}}$ | | | |
| $R_{\text{poly}}$ | 0.77 | | |
| TMT | -0.59 | -0.56 |  |
| AER | -0.58 | -0.46 | 0.46 |
| **SZ** | $R_{\text{mono}}$ | $R_{\text{poly}}$ | TMT |
| $R_{\text{mono}}$ | | | |
| $R_{\text{poly}}$ | 0.37 | | |
| TMT | 0.14 | -0.14 |  |
| AER | - | -0.14 | 0.54 |

### Correlations withclinical symptoms

Task performance was compared with positive and negative symptom scores (SANS and SAPS). Polytonic rhythm showed robust relationships with negative symptoms, a higher aperiodicity detection threshold on this task correlating with the Alogia subscale of the SANS (r_29_=0.38, p<0.04), as well as the SANS total scores (r_28_=0.37,p<0.05), meaning that the greaterthe level of negative symptoms, the larger the additional gap across an inter-tone interval is needed to detect the aperiodicity. Regarding the relationship between performance on the four experimental tasks, and the total and subscale scores for the SAPS and SANS, no other correlations were significant.Finally, to assessputative effects of medication on task performance,and given that the medication regimes for our sample were somewhat heterogeneous, we tested for correlations between task performance and antipsychotic dose (CPZ, chlorpromazine equivalents), but none were significant (all p’s> 0.13).

## DISCUSSION

We initiated this study based on similar demonstrations of pitch correlated prosody deficitsinamusia and schizophrenia. In amusia, rhythm perception abilities had been shown to be pitch dependent. As we noted in the introduction, this finding is at face value counterintuitive: Less pitch sensitivity should if anything mean less influence of tone pitch heterogeneity. Additionally, previous lesion data has demonstrated doubly dissociated neuroanatomical loci for elemental time and pitch processing.These prior pieces of evidenceled to a classical neuropsychological hypothesis that pitch and rhythm are yoked at a higher level – perhaps for their combined use in a language processing circuit likely encompassing Broca’s right homologue (**Figure 3**)(31;32).

**Figure 3.**
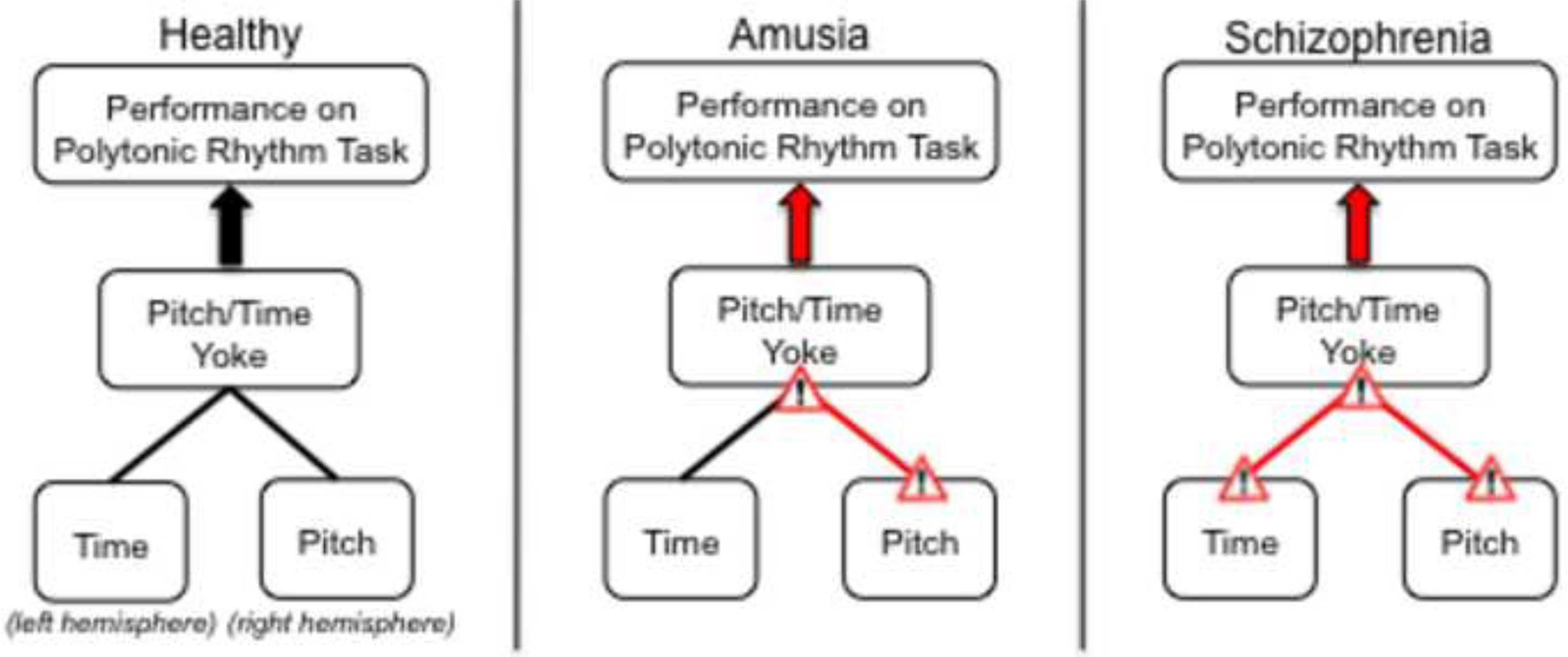
Summary of the pattern of rhythmic dysfunction in healthy, amusia, and schizophrenia populations. **(A)**Hyde & Peretz (2004) and Foxton (2006) posited the following model of rhythm and pitch perception: Perceiving polytonic rhythms integrates information from both time and pitch. **(B)**In amusia, pitch perception deficits result in polytonic rhythm deficits despite intact temporal perception as evidenced by monotonic rhythm performance. **(C)**In schizophrenia, both time and pitch perception deficits again result in impaired polytonic rhythm perception.

Incorporating our schizophrenia data into this classic model suggests that there are deficits in time perception as evidenced by monotonic rhythmperformance, pitch deficits as evidenced by tone matchingperformance, and secondary yoked time/pitch deficits as evidenced by the effects of pitch perception ability on monotonic and polytonic rhythmthreshold differences. The effect of pitch ability on monotonic versus polytonic rhythm perception was observed across control and schizophreniaparticipants, yet the patients with schizophrenia predominated in the low and intermediate task performance groups and control participantsformedthe majority for the high pitch group. Thus, schizophrenia patients displayed time and pitch yoked rhythm deficits, consistent with amusia, yet they displayed impaired pure meter perception, which is unlike amusia.

This pattern of multi-stage perceptual dysfunction is consistent with a narrative in schizophrenia research in which widespread auditorysensory dysfunction, such as time-based duration mismatch negativies, intensity, and pitch, are much in evidence(see(23) for a review).While broad in nature, these inter-related deficits yield a definite structural and hierarchical pattern: for example, low-level automatic recognition of pitch change contributes to independent higher leveldeficits in the detection of pitch deviant targets(22).

Clinically, rhythmic abilities correlated with SANS negative symptoms,but not with positive symptoms. This is consistent with prior research relating auditory processing abnormalities and negative symptoms. Yet, unlike prior data, pitch and prosody task performance did not correlate with negative symptoms. This is the first time such a correlation was not observed, despite multiple replications(2;3;7;41).

In examining schizophreniausing amusiabased rhythm tasks, we gain an important glimpse ofthe structure of auditorysensory processing deficits and how they might vertically integrate in producing vocal social communication dysfunction. The first thing we noticed in our sample was that comparing how performance on polytonic and monotonic rhythm tasks diverges as a function of pitch with how it diverges as a function of prosodic task performance, yielded striking similarities (**Figure1B &D**).This finding, coupled with similar patterns of pitch-prosodytask correlations in the healthy and schizophrenia groups, suggested that diagnosis as an explanatory factor in modeling prosodic dysfunction would exert itself primarily in the degree or manner in which theirimpaired pitch and rhythmic abilities contribute to prosodic dysfunction. To our mind, this is not surprising, as patients display wide ranging task performance, and it likely reflects the heterogeneous nature of the schizophrenia diagnosis in which a subset might display impaired pitch perception while others might not(41).

Inthe schizophrenia and control groups,pitch perception correlated with prosody perception, as has been demonstrated previously(2;3;7;41). Both people with amusia and patients with schizophrenia have pitch deficits, as well as distinctpolytonic rhythmic impairments, hypothetically reflectingaudiolinguistic processing dysfunction at a stage in which pitch and time information converge (see **Figure 3**). Schizophreniaparticipants additionally displayed reductions in pure meter (monotonic rhythm) perception. However, how these deficits contribute to prosodic processing in our participants is unknown, as correlation and regression analyses provided an incomplete picture. Hence, we formallytested a model, which posited both direct and indirect pathways in which time and pitch cues mightcontribute to impaired prosodic processing. The statistically optimal model that we arrived at indicated that elemental pitch perception, as estimated by tonematching, contributed directly to prosodic processing, whereas rhythmic contributions were more nuanced: monotonic rhythmic abilities contribute indirectly to prosodic processing, via contributions to polytonic rhythmic perception. Polytonic rhythmic perception, theoretically indexing yoked time-pitch processing,contributes to prosodic processing dysfunction. This model, while robust and equivalent across all participants and withincontrolsalone, was notably different in schizophrenia,which lacked significant covariance between pure meter perception and tone matching abilities and the magnitude of contribution to prosodic processing from polytonic rhythm processing was weaker.

Differencesobserved in the schizophrenia model versus those across groups and with HC alone need further replication. In schizophrenia, the absence of pitch and monotonic rhythmic covariance is robust (Controls: *ρ*_*s*_=-0.72(0.09), vs. schizophrenia:*p*_*s*_-0.06(0.20)),likely reflecting a true difference, while slighter differences where variance in high, such as a reduced polytonic rhythmprosody path to prosody (schizophrenia: *p*_*s*_=-0.17(0.18) vs. controls*p*_*s*_=0.39(0.18)), need further replication.Neither can we exclude whether the deficits reflect the inattention or global cognitive decline associated with schizophrenia without neuropsychological data, although it might be difficult to imagine that such impairments would have the specific pattern of effects observed on the magnitude of relationship between pitch, rhythm, and prosodic processing. We look forward to remedying theselimitations, as well as the current lack of direct comparative data with amusia participants, in a larger-scale comparative study. A second fruitful avenue for future study is the examination of phonemic processing and phonology more generally in both amusia and schizophrenia. Phonemic distinctions such as /Ba/ versus /Wa/ rely on both changes in intensity of over time and frequency (formant change) over time. Consequently, it will be interesting to assess if putative time pitch or yoked time-pitch deficits in either illness upward generalize to phonemic or phonological dysfunction.

While this study focusedon what schizophrenia researchers can learn from studies of amusia,and how core rhythmic and spectral (i.e. pitch) abilities contribute to prosodic processing more generally, it also offers a more general proof-of-concept for a new approach to the cognitive neuroscience of neuropsychiatric illnesses.Illnesses like schizophrenia are increasingly appreciated as being heterogeneous innature,and abnormal behavior such as prosody perception may be impaired in some individuals, but not in others. Additionally, many of the socio-cognitive markers such as impaired prosody perception that present in schizophrenia are not specific to the illness, and alsopresent in other disorders such as autism, Alzheimer’s disease, bipolar disorder, and Parkinson’s disease(42–46). From a neuroscience perspective, the ability to perceive social intent through prosody is a highly complex event involvingmany cortical and subcortical brain regions (47–50). It is therefore probable that prosodic processing deficits can arise from dysfunction within any one or combination of these systems. Thus, reduced abilities in emotional prosody in differing diagnostic groups that are at *prima facie*similar in terms of dysfunction magnitude,may be generated via aberrations in differing sub-processes or neural mechanisms upon which prosodic processing is built. This approach may be one avenue to examine the Research Domain Criteria (RDOC) proposed by Thomas Insel which emphasizes the exploration of dimensional variation in brain-behavior phenotypes, within and across disorders(51).

Here we “carved prosodic processing at its rhythmic and pitch joints”,and modeled the patterns of interrelationship, dependencies and contributions to prosody perception. This is an approach that can easily be generalized, in the following two ways. Dysprosodia’s ubiquity across multiple disorders is a taxonomic liability when examined alone. However, characterizing prosodic processing as the function of the covariances and number and magnitude of paths between sub-processes like pitch and time,transforms this minus into a plus: Namely, it provides highly sensitive way to differentiate neuropsychiatric illnesses that have dysprosodia as a common trait.These psychophysical “connectomes”(52) would detail the origins of functional deficits and the pathways throughwhich they interact. In this way, the unique pathophysiological mechanisms in schizophrenia could be characterized in relation to other disorders with similar deficits. In addition todisorder-specific connectome paths, the same constructs could be used to design individual-level maps that extend beyond categorical models of mental illness, allowing individualized modeling of cognitive-neurological and auditory-linguistic processing dimensions.

## Acknowledgments

The authors wish to acknowledge Jessica Foxton who kindly lent us therhythm tasksfrom which we derivedour own version, and Jan Richard for her help in programming the rhythm tasks. The authors would also like to thank Monica Calkins and Raquel Gur of the Schizophrenia Research Centerat the University of Pennsylvania, for the their help with the clinical and diagnostic assessment; Sevda A. Zerbe for assistance in data collection, and finally Daniel Wolf for his suggestions on the manuscript.This work was supported in part by NIMH grant K01-094689, NIMH grant T32 MH19112-24, and a young investigator award from the Brain and Behavior Research Foundation (formerly NARSAD).

